# MicroRNA-383-5p alleviates hemorrhagic transformation and improves outcomes after endovascular recanalization for acute ischemic stroke

**DOI:** 10.64898/2025.12.27.696665

**Authors:** Qiu-Yi Jiang, Wang Qin, Jin-Kun Zhuang, Ze-Xin Chen, Qian-Yu Chen, Chang-Luo Li, Zhong-Song Shi

## Abstract

**Background:** Hemorrhage transformation (HT) after reperfusion therapy is linked with poor outcomes in acute ischemic stroke patients. This study aimed to determine the role of neuronal microRNA-383 in alleviating HT-associated injury using middle cerebral artery occlusion (MCAO) and oxygen-glucose deprivation reoxygenation (OGD/R) models.

**Methods:** In neurons following OGD/R, the family of miR-383 and NADPH oxidase (NOX) expression was evaluated. After miR-383-5p intervention and small interfering RNA, reactive oxygen species (ROS) and neuronal injury were assessed. Two hundred and five hyperglycemic rats were used to establish the HT model induced by mechanical reperfusion after 5-hour MCAO, followed by 3 and 6 hours of reperfusion, and treated with intravenous administration of miR-383-5p agomir before reperfusion. Brain water content, hemorrhage severity, infarct volume, blood-brain barrier (BBB) disruption, neuronal apoptosis, ROS production, miR-383-5p and NOX4 expression, and BBB-associated proteins were assessed.

**Results:** MiR-383-5p expression significantly decreased in OGD/R-treated neurons, while miR-383-3p expression did not differ. Elevating miR-383-5p levels reduced neuronal injury, ROS overproduction, and NOX4 overexpression as a target of miR-383-5p, shown in OGD/R-treated neurons. In the MCAO model, increased miR-383-5p suppressed NOX4 upregulation, alleviated brain edema, infarct volume, and hemorrhage severity, reduced neuronal apoptosis and ROS overproduction, and preserved BBB integrity after mechanical reperfusion for ischemia, thereby improving short-term neurological outcomes.

**Conclusions:** MiR-383-5p alleviates oxidative stress injury and neuronal apoptosis, and preserves BBB integrity by regulating NOX4. Neuronal miR-383-5p could become a potential target for intervention to decrease HT and ameliorate outcomes in endovascular reperfusion treatment for acute ischemic stroke.

## Introduction

Reperfusion therapy with thrombolysis and endovascular thrombectomy can benefit acute ischemic stroke (AIS) patients; however, hemorrhagic transformation (HT) following these treatments is associated with poor long-term prognosis.^1,2^ Recent studies have shown that asymptomatic HT, which was previously considered a benign phenomenon with the occurrence in one to three patients after endovascular treatment, can also significantly impact functional outcomes.^3,4^ In light of this new understanding, developing a neuroprotective strategy as an effective adjunct to reperfusion treatment is crucial for alleviating HT, thereby improving outcomes in AIS.

The mechanisms of oxidative stress, immune inflammation, and apoptosis are crucial for understanding reperfusion injury and HT after treatment.^2^ In the central nervous system, nicotinamide adenine dinucleotide phosphate oxidases (NOXs) are the primary sources of reactive oxygen species (ROS) generation, which play an important role in regulating the redox state of neurovascular cell homeostasis and the processes related to stroke.^5^ Targeting NOX4, which is one of the prevalent NOXs in neurons and endothelial cells, can reduce cerebral ischemia-reperfusion injury and minimize hemorrhage by alleviating oxidative stress injury and blood-brain barrier (BBB) disruption during AIS.^6–8^

MicroRNAs have been recognized as potential biomarkers for diagnosis and prognostic prediction in acute ischemic stroke.^9^ MicroRNA-383 is one of the enriched microRNAs in neurons within the brain.^10^ The downregulation of microRNA-383 in clinical blood samples may be associated with HT in AIS patients.^11^ Moreover, decreased levels of microRNA-383 have been observed in peri-infarction tissue of the middle cerebral artery occlusion (MCAO) model with HT induced by mechanical reperfusion.^12^ The protective role of microRNA-383 in HT is yet unclear. Our study aims to investigate the role of neuronal microRNA-383 in the regulation of HT, particularly focusing on NOX signaling mechanisms, through oxygen-glucose deprivation/reoxygenation (OGD/R) conditions in neurons and using the MCAO model with HT induced by mechanical reperfusion in hyperglycemic rats. The goal of this study is to identify a novel therapeutic target to alleviate hemorrhagic injury following endovascular reperfusion treatment.

## Materials and Methods

The study protocol was approved by the Institutional Animal Care and Use Committee in our institution.

### In-vitro studies in the neuron model of OGD/R

Primary mouse cortical neurons were cultured using the previous method.^6, 12^ In OGD/R experiments, the third-generation neurons were deprived of oxygen and glucose in a hypoxic chamber (Stemcell) for 2 hours. Then, neurons were reoxygenated in a normoxic chamber with a glucose-containing medium at dynamic time points from 0 to 72 hours. The expression of miR-383-3p, miR-383-5p, and mRNA of NOX1, NOX2, NOX3, NOX4, Toll-like receptor (TLR) 2, TLR4, Caspase3, Caspase8, and Caspase9 was measured by RT-PCR.

MicroRNA mimics and inhibitors, small interfering RNA (siRNA), and Lipofectamine 3000 (Invitrogen) were used for transfection. MiR-383-5p mimics and inhibitors were used at 50nM and 100nM, respectively. Neurons were incubated with miR-383-5p mimics, miR-383-5p inhibitors, or siRNA NOX4 for 48 hours, then underwent 2-h OGD with reoxygenation at 24 hours. Finally, neurons were collected for RNA and protein assay, ROS measurement, and lactate dehydrogenase assay. MiR-383-5p mimics, miR-383-5p inhibitors, and siRNA targeting NOX4 were provided by RiboBio.

### Rat MCAO developed hemorrhage transformation by mechanical recanalization

We utilized the intraluminal filament technique to occlude the left MCA for 5 hours in adult male Sprague-Dawley hyperglycemic rats (250-280g) to induce severe cerebral ischemia. To create more obvious HT, rats were induced into acute hyperglycemia via multiple intraperitoneal injections of 50% dextrose, as previously reported.^7,13,14^ The rats received reperfusion at 6 hours after a 5-hour occlusion, with retrieval of the intraluminal filament. All experimental procedures were performed under anesthesia using 5% isoflurane for induction and 2% isoflurane for maintenance.

First, 30 hyperglycemic rats were randomly allocated to evaluate miR-383 levels in blood and brain tissue across the following groups: sham operation and 5-hour MCAO with various reperfusion time points (0, 1, 3, and 6 hours). Second, 175 hyperglycemic rats were randomly allocated into five groups to elucidate the neuroprotective role of miR-383-5p intervention: sham operation, MCAO, MCAO with microRNA control intervention, and MCAO receiving miR-383-5p agomir treatment following reperfusion at 3 hours or 6 hours. The miR-383-5p agomir or microRNA control (50 nmol/kg body weight; Gene Pharma) was injected intravenously half an hour before retrieval of the intraluminal filament. After 3 or 6 hours of reperfusion, rats were sacrificed to harvest brain tissue and whole peripheral blood. For RNA analysis of blood, we stored whole blood in PAXgene Blood RNA tubes from PreAnalytiX GmbH. Peri-infarct tissues were collected for RNA and protein assays. Brain hemispheres were collected in OCT compound and snap-frozen. Immunohistochemical staining, TUNEL staining, and ROS measurement were performed with the 7-µm-thick cryosections.

### Assessment of neurological function, infarct volume, BBB permeability, and HT in the rat stroke model

Rats were examined and scored for neurological deficits on a scale from 0 (no neurological deficits) to 4 (absence of spontaneous movement and unconsciousness) at 3 and 6 hours after reperfusion. We utilized 2,3,5-triphenyltetrazolium chloride (TTC) staining and the wet-dry weight method to evaluate infarct volumes and brain edema, respectively. Tail vein injection of Evans blue (EB) solution was performed 3 hours before sacrifice. The absorbance intensity of EB extravasation in each hemisphere was measured to calculate the EB extravasation index, which reflected BBB permeability. Hemoglobin content in each hemisphere was assessed using a spectrophotometric assay to quantify the amount of cerebral hemorrhage. These methods were carried out as previously described ^7, 14^ and provided in the supplementary method for details.

### Quantitative RT-PCR

Quantitative real-time PCR was performed to evaluate miR-383 and gene mRNAs in neurons following OGD/R, as well as in peri-infarct tissue and peripheral blood from the animal model. RNA was isolated using the TRIzol reagent, and primers for all targets were sourced from RiboBio. For normalization, U6 served as the internal reference for miR-383, while β-actin was used for mRNA targets. The expression of miR-383 and mRNAs was quantified using the 2-ΔΔCT method.

### Western blot

The total protein of the neurons following OGD/R and the peri-infarction tissue of the MCAO model were extracted. The levels of NOX4, claudin5, and occludin were assessed by enhanced chemiluminescence. The antibodies to NOX4, claudin5, occludin, and β-actin were used at 1:1000 and were purchased from Abcam. A molecular imager system was used to detect the protein’s expression bands. The ImageJ software was then used to quantify optical density.

### Immunohistochemical fluorescent staining

The brain cryosections were stained using immunohistochemistry. The primary antibodies to NOX4 (Abcam), NeuN as the neuron marker (Millipore), and GFAP as the astrocyte marker (Millipore) were used at 1:200. The fluorescent secondary antibodies Alexa Fluor 594 (Invitrogen) or Alexa Fluor 488 (Invitrogen) were probed for sections at 1:250. A confocal microscope was used to capture three random images from the peri-infarct region in the sections. The ImageJ software was then used to quantify the area of positive fluorescence.

### Reactive oxygen species

ROS in brain cryosections and in an OGD/R neuronal model were examined using DHE (Sigma, D7008) and DCFH-DA (Sigma, D6883) to detect superoxide anion and hydrogen peroxide, respectively. Neurons in the OGD/R model and brain cryosections were incubated with 10 µmol/L of DHE or DCFH-DA for 30 minutes according to the manufacturer’s protocol. The cells and brain sections were photographed under a fluorescence microscope. The ImageJ software was used to quantify the positive fluorescence area for the ROS (superoxide anion and hydrogen peroxide) in the neuronal OGD/R model and in the peri-infarct region of the MACO model.

### Terminal Deoxynucleotidyl Transferase dUTP Nick End Labeling assay

TUNEL and NeuN double staining was performed to assess the neuronal apoptosis in the peri-infarct region, according to the protocol, using the TUNEL in situ cell death detection kit (Roche). The ratio of TUNEL-positive neurons to total neurons was used as the apoptotic index to assess the extent of neuronal apoptosis.

### Lactate dehydrogenase (LDH) assay

LDH release in the neuronal OGD/R model was performed using the LDH Cytotoxicity Colorimetric Assay Kit. LDH activity was assessed by measuring absorbance at 490 nm in each group to evaluate neuronal damage.

### Dual-luciferase reporter assay

Dual-luciferase activity technology was used to clarify the relationship between miR-383-5p and NOX4. The details are provided in the supplementary method.

### Statistical analysis

All statistical evaluations were conducted using SPSS 31.0. Data are expressed as mean ± standard deviation and analyzed using one-way ANOVA or Mann–Whitney U test based on Shapiro–Wilk normality results. A p-value < 0.05 was considered statistically significant.

## Results

### MiR-383-5p inhibited NOX4 overexpression and alleviated neuronal injury following OGD/R

Decreased miR-383-5p expression was shown in neurons following OGD/R, while there was increased expression of NOX1, NOX2, NOX4, TLR2, TLR4, Caspase3, Caspase8, and Caspase9. The expression of miR-383-3p had no significant change after OGDR (Supplementary Figure S1). After intervention with miR-383-5p mimics, NOX4 and TLR4 upregulation in neurons was reduced, whereas the expression of other genes was not altered after OGD/R. MiR-383-5p inhibitor intervention intensified NOX4 upregulation (Figure 1A). NOX4 siRNA, combined with miR-383-5p inhibitor, relieved the NOX4 upregulation, suggesting that miR-383-5p regulates NOX4 in OGD/R-treated neurons (Figure 1B and 1C). Dual luciferase reporter assays, combined with bioinformatic predictions, confirmed that NOX4 is a direct target of miR-383-5p (Figure 1D). LDH release from OGD/R-treated neurons was inhibited by miR-383-5p mimics, whereas worsened by miR-383-5p inhibitor. The influence of miR-383-5p mimics on alleviating neuronal injury is similar to that of the early reperfusion in the OGD/R model with 0.5 hours of ischemia (Figure 1E).

**Figure 1.**
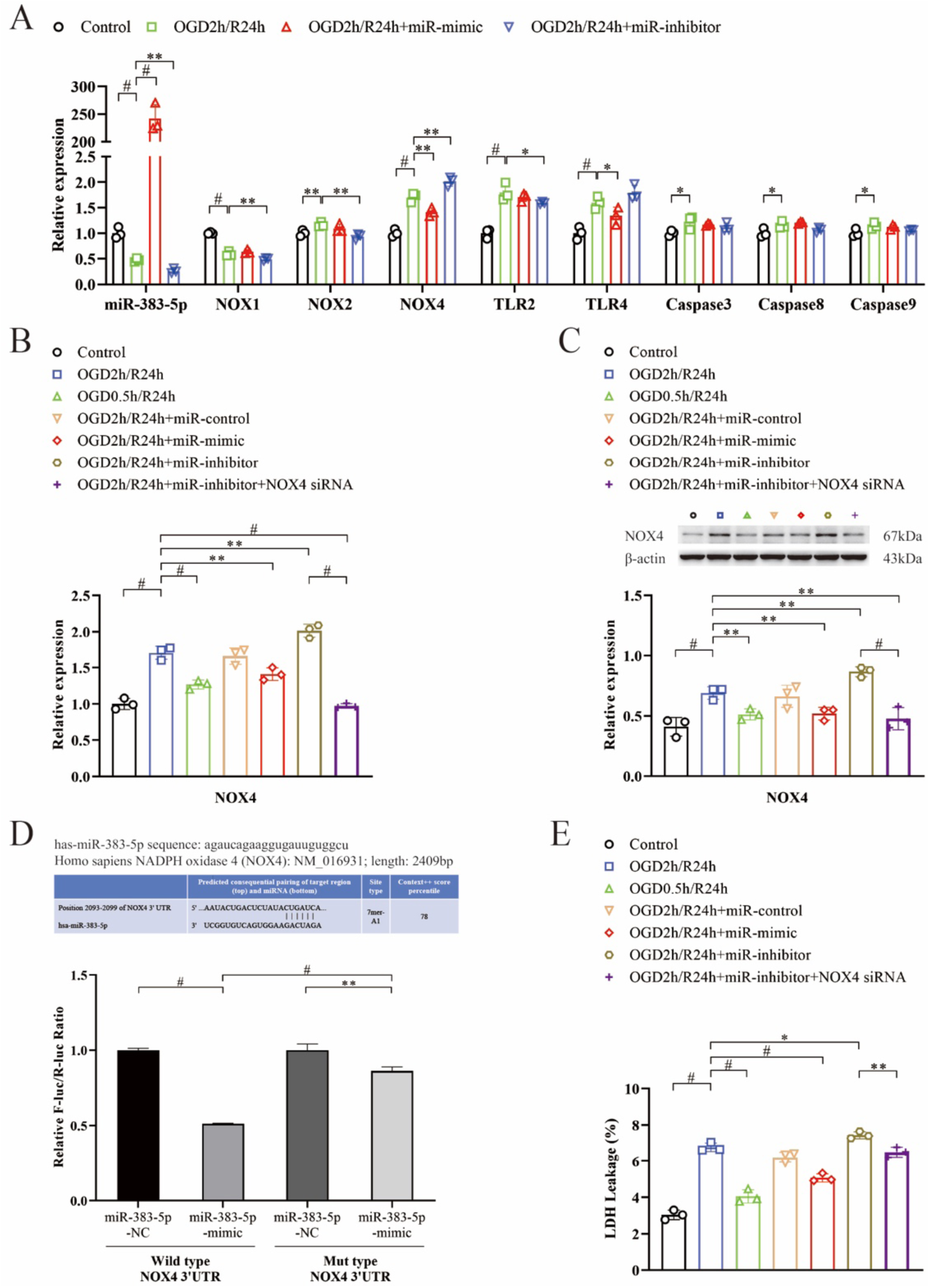
Elevating miR-383-5p levels has a neuroprotective role in the OGD/R model of neurons by suppressing NOX4 overexpression. **A**, miR-383-5p mimics intervention increased miR-383-5p expression and decreased NOX4 upregulation. MiR-383-5p inhibitor intensified NOX4 upregulation. The result of neurons receiving OGD/R with microRNA control intervention was not included in the figure. **B and C**, MiR-383-5p inhibitor intensified NOX4 upregulation. MiR-383-5p inhibitor combined with NOX4 siRNA reversed NOX4 upregulation, as shown by RT-PCR (B) and Western blot (C). **D**, NOX4 is the direct target gene of miR-383-5p determined by a dual-luciferase reporter assay. **E**, miR-383-5p mimics inhibited LDH release from neurons. Each sample was tested in triplicate. NOX, NADPH oxidase; TLR, Toll-like receptor. *P<0.05; **P<0.01; ^#^P<0.001.

### MiR-383-5p inhibited ROS by regulating NOX4 in the neuron model of OGD/R

MiR-383-5p mimic intervention inhibited the rise of ROS (hydrogen peroxide and superoxide anion) in neurons induced by OGD/R. MiR-383-5p inhibitor intensified ROS overproduction following OGD/R. NOX4 siRNA combined with the miR-383-5p inhibitor attenuated ROS upregulation, suggesting that miR-383-5p directly regulates NOX4 to attenuate ROS overproduction (Figure 2).

**Figure 2.**
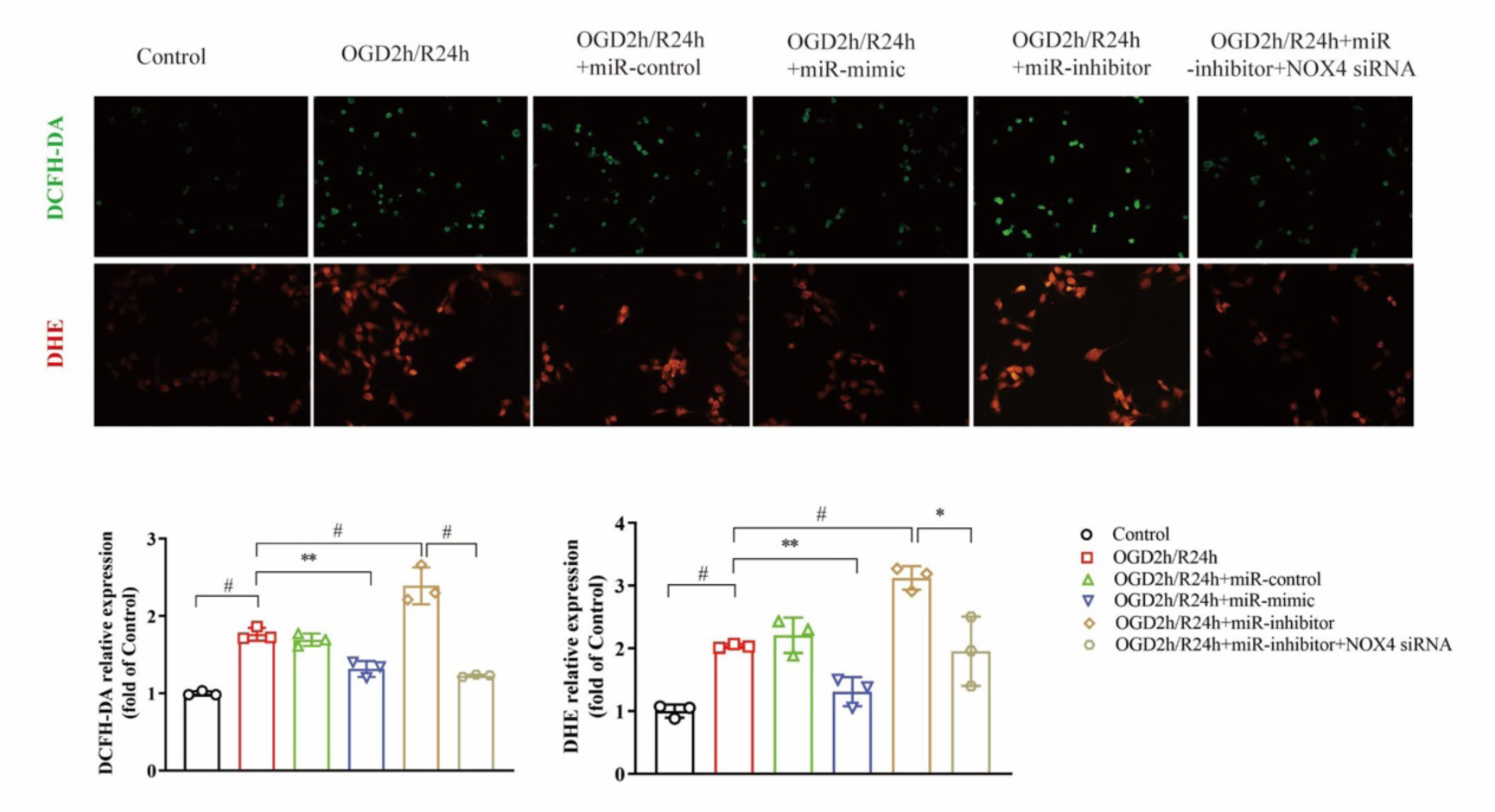
Increased miR-383-5p reduces reactive oxygen species overproduction by suppressing NOX4 in the neuronal OGD/R model. Detection of hydrogen peroxide by DCFH-DA and superoxide anion by DHE in neuronal OGD/R model treated with miR-383-5p mimic and inhibitor, and NOX4 siRNA. Each sample was tested in triplicate. *P<0.05; **P<0.01; ^#^P<0.001.

### MiR-383-5p reduced NOX4 overexpression, ROS overproduction, and neuronal apoptosis in the MCAO model

In the MACO rats, miR-383-5p was downregulated in the peri-infarction and blood after reperfusion at 1, 3, and 6 hours (Figure 3A). Upregulated miR-383-5p levels occurred in both blood and peri-infarction after intravenous injection of miR-383-5p agomir compared with the ischemia/reperfusion group (Figure 3B and Supplementary Figure S2). The expression of NOX4 in peri-infarction tissue was increased in the ischemia-reperfusion group, whereas miR-383-5p agomir intervention suppressed this increase (Figure 3B). The patterns of NOX4 expression changes occurred in neurons within the peri-infarction region, and also showed in astrocytes within the peri-infarction region (Figure 3C). ROS overproduction in the peri-infarction region was revealed by DCFH-DA and DHE staining after ischemia-reperfusion, whereas miR-383-5p agomir significantly attenuated ROS overproduction (Figure 3D). TUNEL-positive neurons reflecting apoptosis in the peri-infarction were much more increased after ischemia/reperfusion, but reduced expression after miR-383-5p agomir treatment (Figure 3E).

**Figure 3.**
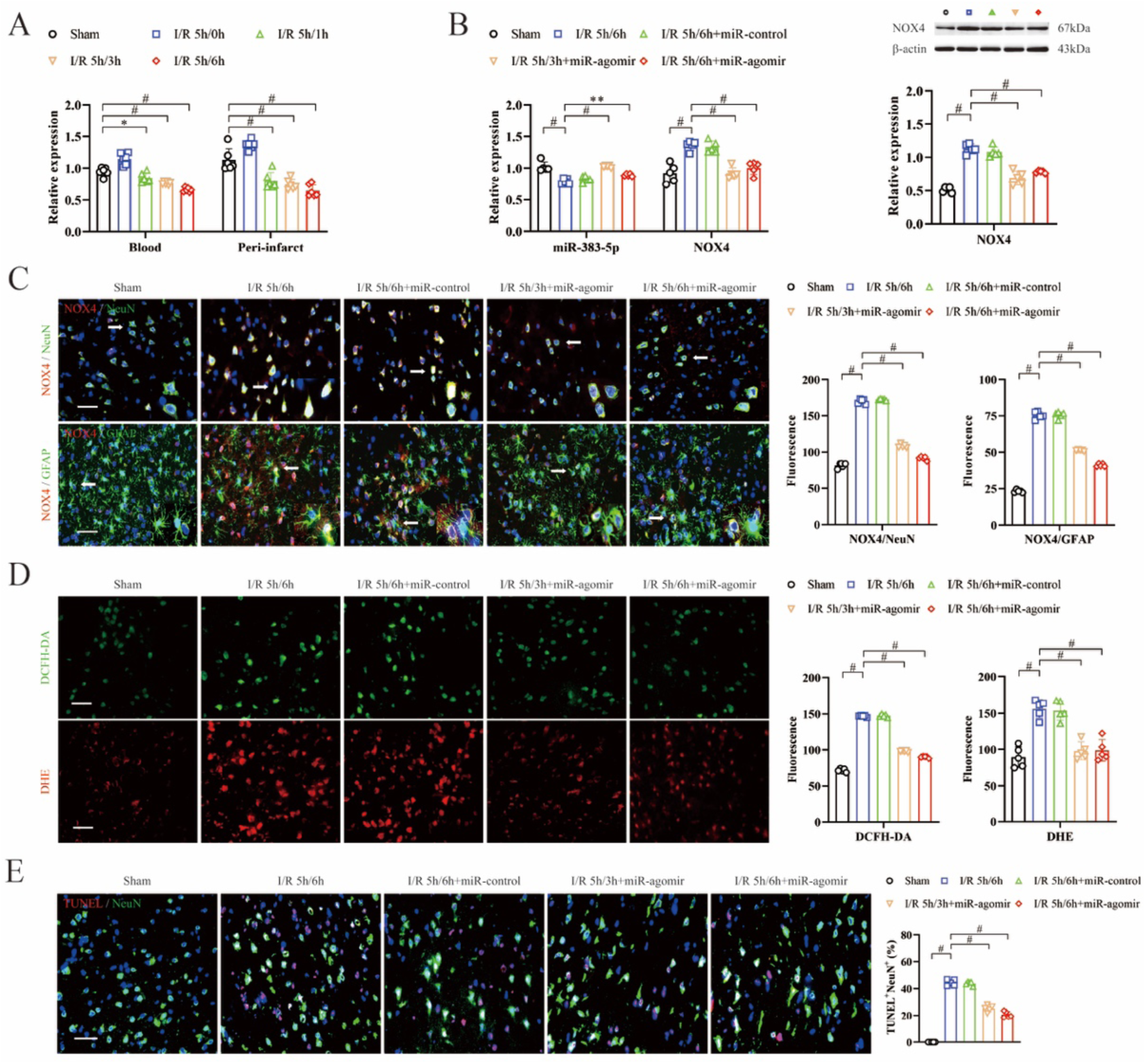
MiR-383-5p agomir protects the brain peri-infarction against oxidative damage and apoptosis by regulating the NOX4/ROS signaling pathway in a hyperglycemia-associated reperfusion-induced hemorrhagic transformation model. **A,** miR-383-5p expression decreased in the blood and peri-infarction tissue of the MCAO model (n=6). **B,** miR-383-5p agomir treatment increased the miR-383-5p expression and reduced NOX4 expression in the peri-infarction tissue of the MCAO model (left, RT-PCR; right, Western blot) (n=5). **C,** miR-383-5p agomir treatment reduced NOX4 expression in neurons and astrocytes in peri-infarction tissue, as shown by immunofluorescence staining (n=5). **D,** miR-383-5p agomir treatment reduced ROS overproduction in the peri-infarction region, as quantified by DCFH-DA and DHE staining (n=5). **E,** miR-383-5p agomir treatment reduced neuronal apoptosis in the peri-infarction, as shown by TUNEL and NeuN immunolabeling (n=5). Scale bars represent 50 µm. MCAO, middle cerebral artery occlusion; NOX, NADPH oxidase; ROS, reactive oxygen species. *P<0.05; **P<0.01; ^#^P<0.001.

### MiR-383-5p suppressed brain edema, infarct volume, and hemorrhage with improved outcome

MiR-383-5p agomir treatment decreased infarct volume in TTC staining (19.9% versus 33.8%, p <0.001, at 6 hours) (Figure 4 A), and reduced hemorrhage severity shown by hemoglobin content index assessment (1.2 versus 1.5, p =0.026, at 6 hours) (Figure 4B), and alleviated brain water content (Figure 4C), compared with rats receiving ischemia-reperfusion. Rats receiving miR-383-5p agomir showed a lower neurological deficit score than those in the ischemia-reperfusion group (1.1 versus 2.2, p <0.001, at 6 hours).

**Figure 4.**
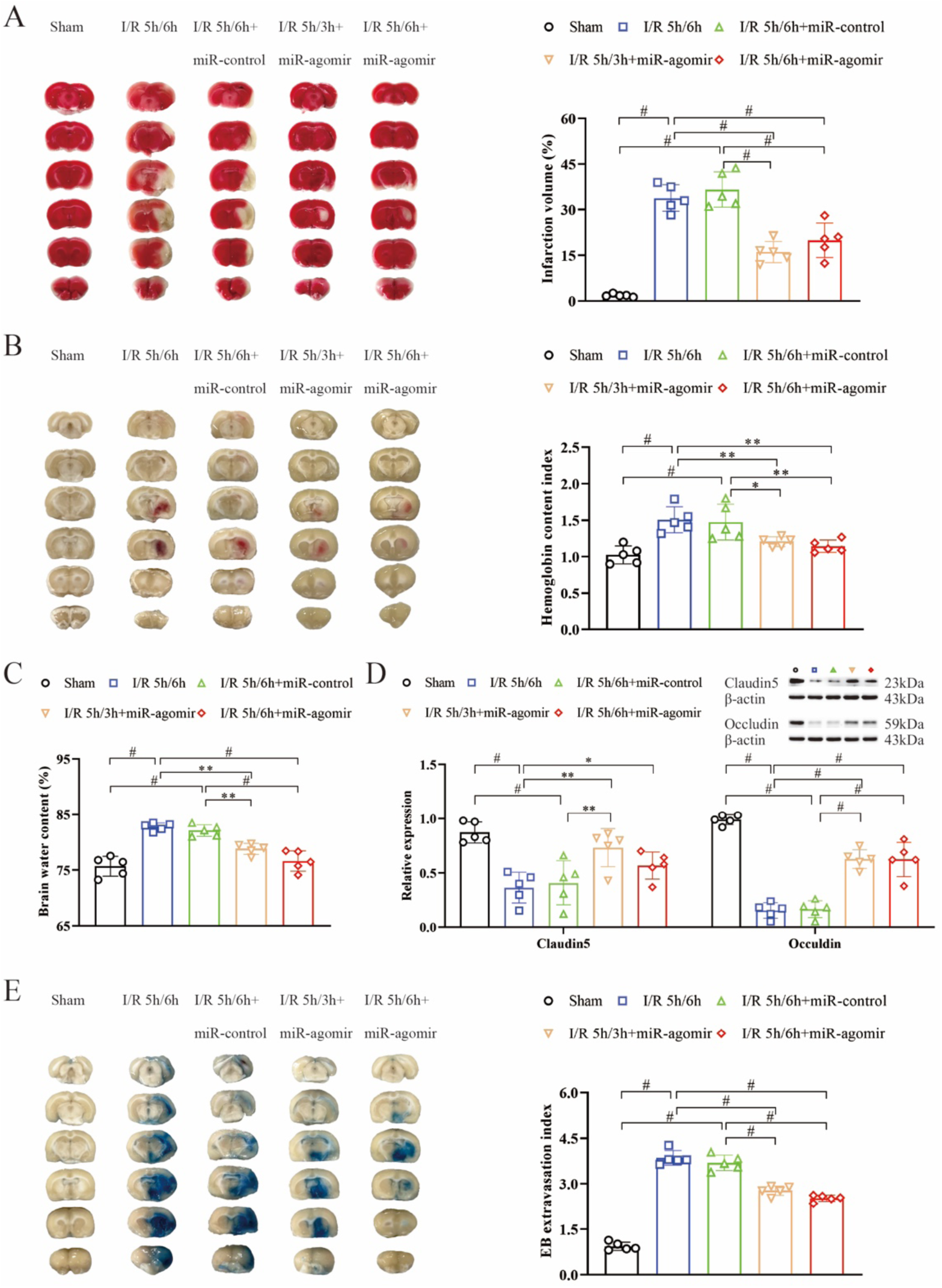
MiR-383-5p agomir reduces infarct volume, hemorrhage, brain edema, and blood-brain barrier disruption. **A**, TTC staining of brain slices and quantification for infarct volume at 3 and 6 h after reperfusion with 5-hour ischemia (n=5). **B**, Hemoglobin content index for assessment of hemorrhage severity (n=5). **C**, Brain water content for assessment of brain edema (n=5). **D**, MiR-383-5p agomir restored the claudin5 and occludin expression in the peri-infarction tissue, as shown by Western blot (n=5). **E**, Blood-brain barrier permeability in the ischemic hemisphere was shown by Evans blue staining (n=5). *P<0.05; **P<0.01; ^#^P<0.001.

### Preservation of the BBB by increased miR-383-5p

MiR-383-5p agomir treatment partly restored the expression of claudin5 and occludin, which were decreased in the peri-infarction tissue after ischemia-reperfusion (Fig. 4D). The marked EB extravasation after ischemia-reperfusion was significantly decreased after miR-383-5p agomir treatment (2.5 versus 3.8, p<0.001, at 6 hours), suggesting that BBB integrity was protected by increased miR-383-5p (Figure 4E).

## Discussion

Our study shows that miR-383-5p downregulation and NOX4 upregulation in the penumbra are associated with hemorrhage after reperfusion intervention in acute ischemic stroke, as demonstrated in the neuronal OGD/R model and the rat MACO model. Increased miR-383-5p expression can inhibit ROS overproduction and neuronal apoptosis by suppressing NOX4 upregulation, a direct target gene, and preserve BBB-associated junction proteins. This process reduces brain edema, infarction volume, hemorrhage, and BBB breakage, then alleviating neurological deficits. These findings provide new evidence that neuronal miR-383-5p/NOX4/ROS signaling is a promising target for reducing reperfusion injury and HT in acute ischemic stroke.

MiR-383, one of the most abundant neuron-associated microRNAs, is associated with various neurological disorders by regulating cell apoptosis, cell proliferation, and oxidative stress. In patients with Alzheimer’s disease, miR-383-5p expression was reduced in blood compared with healthy volunteers. Upregulated miR-383-5p attenuates Aβ-triggered neuronal apoptosis and oxidative stress injury by reducing kinesin family member 1B expression, a direct target gene.^15^ A previous study using the neuronal OGD/R and the MCAO models showed that miR-383-5p protects against ischemic brain injury by regulating histone deacetylase 9 (HDAC9) activity. CCCTC binding factor downregulates miR-383-5p and upregulates HDAC9 expression in cerebral ischemia, then promotes neuronal apoptosis and aggravates neuronal damage induced by endoplasmic reticulum stress.^16^ These findings support our results that increased neuronal miR-383-5p can provide a neuroprotective effect in alleviating cerebral ischemic injury. In our neuronal model study, the miR-383-3p isoform shows no significant change after OGD/R. The isoform −5p and −3p of miR-383 may have a different role in stroke-induced brain injury. Activated microglia by intracerebral hemorrhage transfer miR-383-3p into neurons, which then negatively regulate activating transcription factor 4 expression and promote neuronal necroptosis.^17^ In the microglia OGD/R model, miR-383-3p directly targets the downstream src homology 2 domain-containing phosphatase 2 (SHP2), then regulates nucleotide-binding oligomerization domain-like receptor protein 3 (NLRP3) and its associated cytokines. Transcranial focused ultrasound stimulation exerts a neuroprotective effect in ischemic stroke by suppressing NLRP3 through the Nespas/miR-383-3p/SHP2 pathway.^18^

Increasing neuronal miR-383-5p levels exerts neuroprotective effects by regulating its direct target gene, NOX4, as shown in our results. In humans, seven NOX isoforms produce superoxide anion and hydrogen peroxide to maintain the NOX-ROS signaling balance under normal physiological conditions. In cases of cerebral ischemia and intracerebral hemorrhage, a disruption in redox balance, caused by excessive induction of NOX, initiates a high level of ROS that exacerbates injury to neurovascular cells.^5^ NOX4-centered ROS signaling can be triggered by cerebral ischemia, resulting in oxidative stress damage and neuronal apoptosis. Treatment with an NOX4 isoform-specific inhibitor or genetic inhibition of NOX4 reduces NOX4 overexpression and excessive ROS generation, thereby protecting against ischemia-induced neuronal damage.^6,19^ NOX4 is also involved in oxidative stress injury and neuronal ferroptosis following intracerebral hemorrhage, making it a neuroprotective target for alleviating the hemorrhage burden and secondary brain injury.^20,21^ After cerebral ischemia, neuronal NOX4 contributes to neuronal damage, while endothelial NOX4 disrupts the integrity of the BBB.^22^ MiR-100-5p-loaded exosomes alleviate oxidative stress damage by targeting NOX4 directly in brain endothelial cells OGD/R model, and reduce infarct volume and improve functional outcome in the MCAO model.^8^ The reduced level of miR-383-5p has been previously observed in neuronal and astrocytic models of OGD/R; however, its expression does not change in the OGD/R model of cerebral endothelial cells, pericytes, and microglia cells.^12^ In our study, the beneficial effects on HT associated with increased miR-383-5p may not be directly linked to the inhibition of endothelial NOX4. Excessive ROS generated by NOX4 in neurons and astrocytes, due to the decreased miR-383-5p following ischemia-reperfusion injury, can trigger brain microvascular endothelial barrier dysfunction and BBB disruption by downregulating junctional complexes between endothelial cells and activating matrix metalloprotease. Increased miR-383-5p treated with a mimic inhibits neuronal and astrocytic NOX4 expression and reduces excessive ROS production. Consequently, this results in the restoration of tight junction proteins (claudin5 and occludin) in the peri-infarction region, and a decrease in HT, as observed in our animal study.

The study has several limitations. The effect of increased miR-383-5p on ameliorating long-term outcomes was not explored, and neurobehavioral function assessments were not conducted. Additionally, other target genes directly regulated by miR-383-5p function as neuroprotectors to alleviate hemorrhage after mechanical reperfusion, which warrants further clarification through RNA sequencing in combination with animal studies and neurovascular cell OGD/R models.

## Conclusions

The neuroprotective role of miR-383-5p for ischemia-reperfusion injury has been highlighted in our study. MiR-383-5p alleviates oxidative stress injury and neuronal apoptosis, and preserves BBB integrity by targeting NOX4. Neuronal miR-383-5p could become a potential target for intervention to decrease hemorrhage and ameliorate functional outcomes in endovascular treatment for acute ischemic stroke.

## Author contribution statement

Z-SS participated in the design of the present study. All authors participated in the interpretation and collection of the data. Q-YJ, WQ and Z-SS wrote the initial manuscript. Z-SS revised the manuscript. All authors critically reviewed and edited the manuscript and approved the final version.

## Ethics statements

The study was reviewed and approved by the Institutional Animal Care and Use Committee of Sun Yat-sen Memorial Hospital, Sun Yat-sen University.

